# MGEnrichment: a web application for microglia gene list enrichment analysis

**DOI:** 10.1101/2021.06.10.447854

**Authors:** Justin Jao, Annie Vogel Ciernia

## Abstract

Gene expression analysis is becoming increasingly utilized in neuro-immunology research, and there is a growing need for non-programming scientists to be able to analyze their own genomic data. MGEnrichment is a web application developed both to disseminate to the community our curated database of microglia-relevant gene lists, and to allow non-programming scientists to easily conduct statistical enrichment analysis on their gene expression data. Users can upload their own gene IDs to assess the relevance of their expression data against gene lists from other studies. We include example datasets of differentially expressed genes (DEGs) from human postmortem brain samples from Autism Spectrum Disorder (ASD) and matched controls. We demonstrate how MGEnrichment can be used to expand the interpretations of these DEG lists in terms of regulation of microglial gene expression and provide novel insights into how ASD DEGs may be implicated specifically in microglial development, microbiome responses and relationships to other neuropsychiatric disorders. This tool will be particularly useful for those working in microglia, autism spectrum disorders, and neuro-immune activation research. MGEnrichment is available at https://ciernialab.shinyapps.io/MGEnrichmentApp/ and further online documentation and datasets can be found at https://github.com/ciernialab/MGEnrichmentApp. The app is released under the GNU GPLv3 open source license.

## Introduction

With the recent advances in sequencing technology, researchers are increasingly able to generate larger amounts of genomic data. Investigating changes in gene expression has allowed neuroscientists to move beyond the high-level analysis of cellular dynamics, and into the investigation of the molecular and biochemical pathways and networks underlying brain disorders (1). For example, in the developing brain early life insults can affect rapid and long-lasting changes to gene expression that alter the neuro-immune system and behaviour(2,3).

Microglia, the brain’s resident innate immune cells, appear particularly vulnerable to early life genetic and environmental risk factors for neurodevelopmental, psychiatric and neurodegenerative disorders (4). As sequencing costs have dropped in recent years (6) and the ability to isolate microglial populations from the brain has expanded, a number of key microglial signature gene lists have been identified across disease models (7) and development (8–11). The ease at which this data can be generated and incorporated into various experiments has led to gene expression analysis now being utilized not just in hypothesis testing, but also in hypothesis generation (5). These microglial gene expression differences have been successfully examined across labs and contexts to identify conserved targets and patterns disrupted across brain disorders (2,12). However, there is currently no central repository for published microglial gene lists nor a user friendly, non-programmatic interface that allows biologists to statistically test their gene list of interest for enrichment of identified microglial gene lists from other studies.

Several enrichment tools currently exist to assist users in interrogating their gene expression results, such as enrichment of Gene Ontologies using tools such as DAVID (13), Gene set enrichment analysis (GESA) (14), or pathways KEGG (15,16). However, these interfaces are not specific to individual cell types nor brain disorders and may not accurately reflect microglial-specific processes or disease states. In comparison, direct gene list comparisons to published microglia datasets can lead to cell type or cell state specific insights into underlying microglial mechanisms. However, this requires access to both a curated database of microglial gene lists and the programmatic skills to implement the analysis and statistics. These obstacles can present a daunting challenge for the non-programming wet-lab scientist. With the increasing use of RNAseq and other expression analysis approaches by biologists, there is a growing need for non-programming based tools that allow for efficient analysis without extensive bioinformatic experience. This need is particularly great in the area of neuro-immunology which attracts researchers from a broad set of backgrounds such as neuroscience, immunology, and others.

Our lab has thus developed MGEnrichment (Microglia Enrichment), a customized web application for performing enrichment testing on a manually curated database of gene lists pertinent to microglia. A key feature of our application is the user’s ability to easily upload a list of genes of interest, as well as the accessibility of customizing background gene list settings. The application is intended for use by wet lab scientists who wish to quickly assess the relevance of their gene expression results, and will be of particular interest to those working in the field of microglia research, brain disorders, and neuro-immune activation.

## Design and Implementation

The base functionality of the app was built using the R Shiny package (https://shiny.rstudio.com/), and hosted using shinyapps.io by RStudio. MGEnrichment allows the user to upload a list of genes from their experiment in three common gene identifier (ID) formats (Ensembl, Entrez, Mouse gene symbols (MGI)). Depending on which gene ID format is entered, the database of microglial gene lists (queried from the R biomaRt package (17)) is filtered for the matching ID type. Users can select between setting the background as all mouse genes, all the genes in the microglial gene list database, or an optional user-specified list of background genes. MGEnrichment then performs a one-tailed Fisher’s exact test using the GeneOverlap package (18) to compare the overlap between the user’s input list and each list in the microglia database. Statistical significance is calculated relative to the background gene list and a False Discovery Rate (FDR) correction is then applied across all comparisons. The level of FDR correction is controlled by the user, allowing for greater flexibility in the statistical threshold used for significance determination.

Enrichment results display several key output variables including the odds ratio, p-value, FDR corrected p-values, and the number and IDs of the overlapping genes for each database list.

Information is also provided regarding individual microglial database gene lists including the group they belong to, a description of the gene list, the species the gene list was collected from, as well as a literature source for where the gene list originates. These results may be viewed directly on the web browser, or as a downloaded CSV file.

The database contains 166 unique microglial gene lists from 40 publications pulled from the microglial literature (Supplemental Table 1). Gene lists from mouse, rat and human are included, but all gene IDs were converted to mouse for inclusion in the database. The database of gene lists was manually curated from previous literature using Ensembl IDs, then queried against biomaRt to match the additional corresponding MGI symbols and Entrez IDs. It includes a wide assortment of microglial relevant gene lists collected from multiple treatments, disease states and developmental timepoints in microglia or brain. The default conditions include all genes lists in the database for analysis, but users may also select subsets of gene lists based on six different list categories (groups). Group options include Microglia, Microglia Development, Neuropsychiatric & Neurodevelopmental Disorders human brain, Autism genetics, Autism regulators, and Inflammation. The user can select the groups to be included in the analysis, allowing for more targeted analysis to a specific subgrouping within the database.

To demonstrate the utility of our approach we created two “toy” datasets that examine gene regulation in ASD. Microglial dysregulation has been observed in ASD postmortem brain samples in terms of altered cellular morphology and gene expression. Specifically, there have been four large scale, recent RNAseq studies examining differentially expressed genes from human ASD postmortem brain compared to matched controls (19–22). All four identified immune, and specifically microglial, gene expression as altered in ASD brain (19–22). We took the published gene lists from these papers, divided them into genes with either increased or decreased expression in ASD and then overlapped the four sets to identify genes consistently identified in at least 3 out of the 4 datasets. Gene lists were then converted to mouse Ensembl Identifiers using biomart. Users can access these datasets by clicking their respective buttons on the application and querying the database to look for gene list enrichments. Alternatively, a compiled supplemental excel spreadsheet titled (Supplement Table 2) of both toy datasets and the corresponding MGErichment results can be downloaded from the GitHub repository.

Enrichments were calculated using Ensembl gene IDs, with the background set to “All Genes in the Database”, queried against all gene list groups, and with FDR filtering for q<0.05.

## Results

The MGEnrichment app is setup so that users can easily query the microglia database to analyze the gene expression profiles of their lists compared to selected lists from the database. The provided toy ASD increased gene expression dataset (ASD>CTRL DEGs) produces numerous significant (FDR q<0.05) enrichments with database gene lists. For example, ASD>CTRL DEGs are significantly enriched for genes with increased in expression in schizophrenia, a relationship previously identified(22). There were also significant enrichments with gene lists important for microglial development, gene regulation (*Sall1* and *Mef2c*) and immune activation (PolyI:C and LPS treatments) (Supplemental Table 2). There were also significant enrichments with gene lists generated from microglia from germ free mice, supporting a recent growing literature on the role of the microbiome in ASD (23) and suggesting microbiome disturbances associated with the disorder may contribute to altered brain microglia. From these enrichments, individual genes of interest can be identified among the shared genes to identify novel targets for further investigation. For example, the genes shared by our target toy list (ASD>CTRL DEGs) share several transcription regulators with the microglial lists from germ-free mice, indicating that *Hsbp1, Tgif1*, and *Cebpb* might be reasonable target genes for further exploration.

Similarly, using the ASD decreased gene expression dataset (ASD<CTRL DEGs) produces significant overlaps with lists for other human neuropsychiatric disorders as well as genes regulated in microglial development (Supplemental Table 2). The developmental list enrichments all center around lists of differentially expressed genes between embryonic day 18 (E18) microglia and postnatal microglia (P4, P14 and P60), suggesting that genes with disruption in ASD may impact embryonic microglia maturation towards a postnatal transcriptome.

Together, our two example datasets demonstrate the utility of MGEnrichment in exploring microglial gene regulation in neurodevelopmental disorders. The app can provide both novel insights into differentially expressed gene lists, as well as identification of microglial target genes for further examination.

## Availability and Future Directions

The code for the application is also freely available on our GitHub repository, and released under the GNU General Public License version 3 (GPLv3). By releasing this under an open source license, we aim to provide transparency as to how our program was designed, as well as invite collaboration and contributions from others in the field. Documentation for MGEnrichment is provided within a “help” tab of the web application and at https://github.com/ciernialab/MGEnrichmentApp. All source code is included on the GitHub repository, including the microglia gene list database and instructions for adding in new custom gene lists to the database.

MGEnrichment allows for a targeted approach to understanding microglial biology by leveraging known changes in gene expression across different disease and developmental states. As genomics becomes increasingly intertwined with neuro-immunology and behavioural neuroscience research, the ability to interpret gene expression results within the broader context of microglial biology will be a key skillset for many researchers. We have developed MGEnrichment to accomplish two main goals: firstly, to disseminate an easy to access database of curated microglia-relevant gene lists; secondly, to provide a user-friendly interface for non-programmers to examine their gene lists of interest for impacts on microglial biology.

MGEnrichment’s hosting on the web through the R Shiny platform allows any user to easily query their gene list of interest and download their results for further analysis.

Future directions for the project include expansion to allow for direct comparisons of human gene IDs. We can also expand to include additional types of data visualization, such as dot plots to better visualize the level of gene enrichment and network visualizations to support more systems-based analyses. It is our hope that this app will act as a useful tool to bridge the gap between wet and dry-lab scientists in microglial research, and to help traditional behavioural neuroscientists and immunologists to interpret changes in microglial gene regulation.

## Acknowledgements

We would like to thank members of the Ciernia, Tropini and Osborne labs at UBC for helpful feedback on this project.

## Author Contributions

A.C. conceived of the project, collected the microglia gene list database and contributed to code. J.J. wrote the majority of the code and implemented the R Shiny application. Both authors wrote and edited the manuscript.

## Figure legends

**Figure 1.**
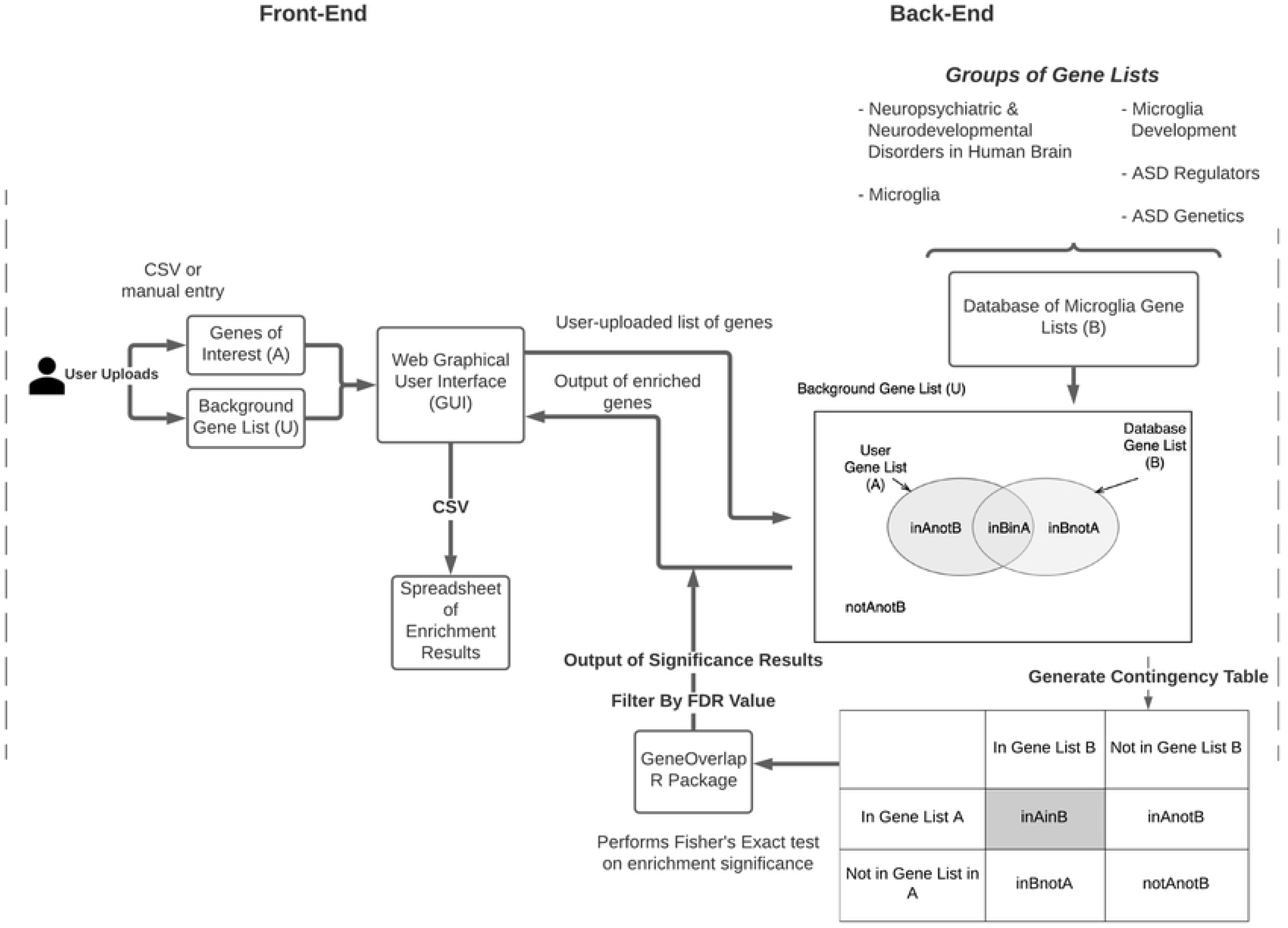
Model of MGEnrichment. Users can upload their gene lists of interest either through a CSV file or through entry into the GUI. The input dataset is compared against the database of microglia gene lists to determine enrichment. The GeneOverlap package is used to calculate a one-tailed Fisher’s Exact Test for enrichment in each gene list, and FDR correct p-values are then calculated across all comparisons. The enriched gene results and corresponding statistical significance are then viewable via the GUI, or exportable via CSV. ASD = Autism Spectrum Disorder.

**Figure 2.**
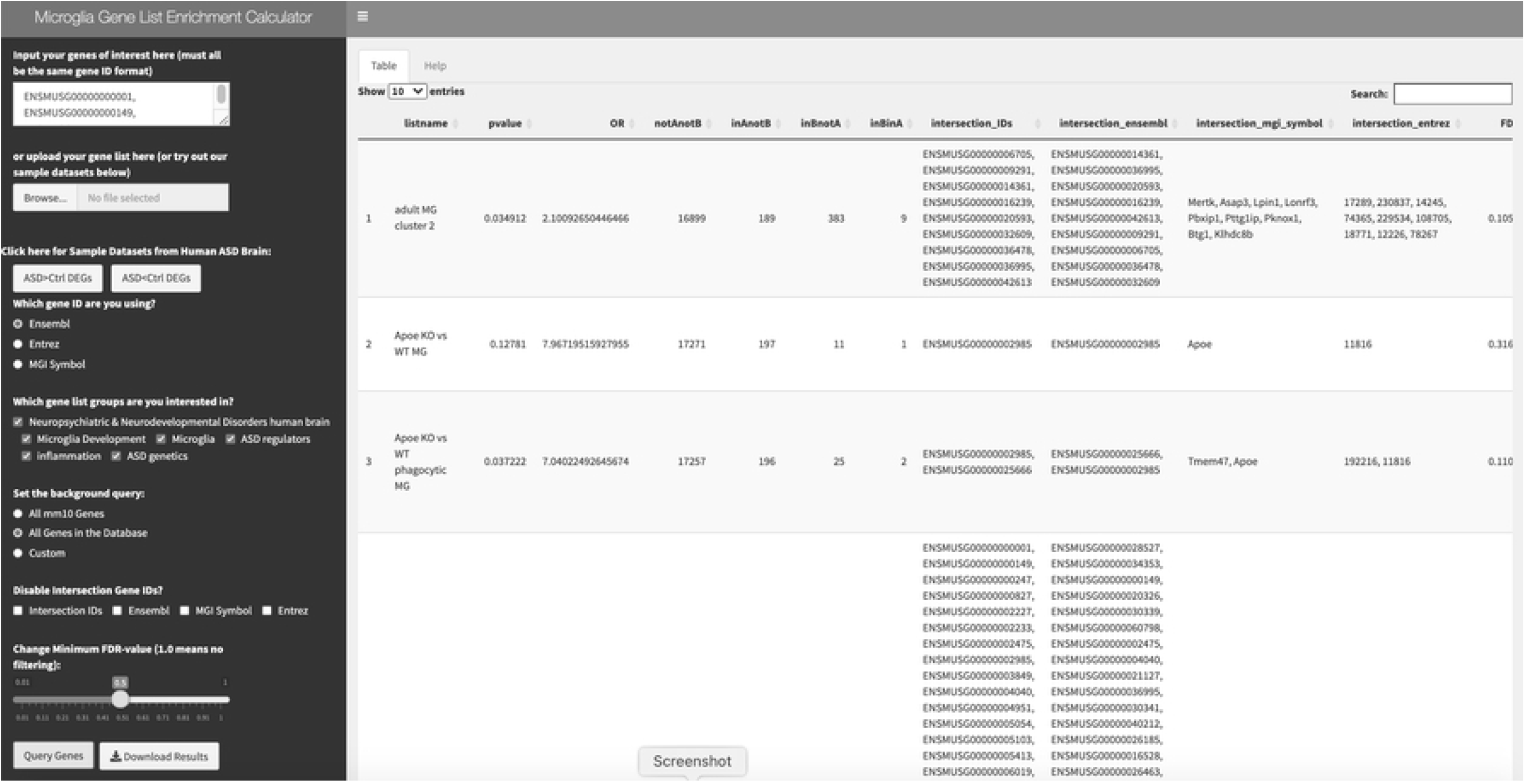
Preview of MGEnrichment, as previewed on a Web Browser. The left panel includes user-input and possible modifications to results, while the table on the right outputs the user query results for each gene list.

## Supplementary Table Captions

### Supplemental Table 2: MG Database

Sheet 1: MG Database. Includes an entry for each gene list in the curated database, description of the gene list, source/citation, group assignment, species of the original study, tissue, abbreviated name and the number of Ensembl mouse IDs within that list.

### Supplemental Table 2: Toy Dataset

Sheet 1: ASD>CTRL_DEGs_Dataset. Includes the input dataset containing the mouse Ensembl IDs for genes identified across 3 out of 4 human brain RNA-seq studies comparing brain samples from ASD and Controls. DEGs show higher expression in ASD compared to Control samples.

Sheet 2: ASD>CTRL_DEGs_Results. FDR filtered (q<0.05) enrichment results are shown for all significant enrichments between ASD>CTRL DEGs and gene lists in the MGEnrichment database

Sheet 3: ASD<CTRL_DEGs_Dataset. Includes the input dataset containing the mouse Ensembl IDs for genes identified across 3 out of 4 human brain RNA-seq studies comparing brain samples from ASD and Controls. DEGs show lower expression in ASD compared to Control samples.

Sheet 4: ASD<CTRL_DEGs_Results. FDR filtered (q<0.05) enrichment results are shown for all significant enrichments between ASD<CTRL DEGs and gene lists in the MGEnrichment database.

## References

1. Geschwind DH, Konopka G. Neuroscience in the era of functional genomics and systems biology. Nature. 2009.

2. Vogel Ciernia A, Careaga M, LaSalle JM, Ashwood P. Microglia from offspring of dams with allergic asthma exhibit epigenomic alterations in genes dysregulated in autism. Glia [Internet]. 2018 Nov 14 [cited 2017 Nov 30];66(3):505–21. Available from: http://doi.wiley.com/10.1002/glia.23261

3. Schwartzer JJ, Careaga M, Coburn MA, Rose DR, Hughes HK, Ashwood P. Behavioral impact of maternal allergic-asthma in two genetically distinct mouse strains. Brain Behav Immun [Internet]. 2017 [cited 2017 Jan 28];36:99–107. Available from: http://www.ncbi.nlm.nih.gov/pubmed/27622677

4. Thion MS, Garel S. On place and time: microglia in embryonic and perinatal brain development [Internet]. Vol. 47, Current Opinion in Neurobiology. 2017 [cited 2018 Mar 22]. p. 121–30. Available from: http://www.ncbi.nlm.nih.gov/pubmed/29080445

5. Holtman IR, Raj DD, Miller JA, Schaafsma W, Yin Z, Brouwer N, et al. Induction of a common microglia gene expression signature by aging and neurodegenerative conditions: a co-expression meta-analysis. Acta Neuropathol Commun [Internet]. 2015 [cited 2017 Jan 24];3(1):1–18. Available from: http://www.actaneurocomms.org/content/3/1/31

6. Wetterstrand K. DNA Sequencing Costs: Data from the NHGRI Genome Sequencing Program (GSP) [Internet]. genome.gov. 2021. Available from: www.genome.gov/sequencingcostsdata.

7. Keren-Shaul H, Spinrad A, Weiner A, Matcovitch-Natan O, Dvir-Szternfeld R, Ulland TK, et al. A Unique Microglia Type Associated with Restricting Development of Alzheimer’s Disease. Cell [Internet]. 2017;169(7):1276-1290.e17. Available from: http://dx.doi.org/10.1016/j.cell.2017.05.018

8. Matcovitch-Natan O, Winter DR, Giladi A, Vargas Aguilar S, Spinrad A, Sarrazin S, et al. Microglia development follows a stepwise program to regulate brain homeostasis. Science. 2016;353(6301):aad8670.

9. Thion MS, Low D, Silvin A, Chen J, Grisel P, Schulte-Schrepping J, et al. Microbiome Influences Prenatal and Adult Microglia in a Sex-Specific Manner. Cell [Internet]. 2018;172(3):500-516.e16. Available from: https://doi.org/10.1016/j.cell.2017.11.042

10. Hammond TR, Dufort C, Dissing-Olesen L, Giera S, Young A, Wysoker A, et al. Single-Cell RNA Sequencing of Microglia throughout the Mouse Lifespan and in the Injured Brain Reveals Complex Cell-State Changes. Immunity. 2019;

11. Li Q, Cheng Z, Zhou L, Darmanis S, Neff NF, Okamoto J, et al. Developmental Heterogeneity of Microglia and Brain Myeloid Cells Revealed by Deep Single-Cell RNA Sequencing. Neuron. 2019;

12. Vogel Ciernia A, Laufer BI, Hwang H, Dunaway KW, Mordaunt CE, Coulson RL, et al. Epigenomic Convergence of Neural-Immune Risk Factors in Neurodevelopmental Disorder Cortex. Cereb Cortex [Internet]. 2019 Jun 26 [cited 2019 Jul 8];bhz115. Available from: http://www.ncbi.nlm.nih.gov/pubmed/31240313

13. Jr GD, Sherman BT, Hosack DA, Yang J. DAVID: Database for Annotation, Visualization, and Integrated Discovery. 2003;4(5):P3.

14. Subramanian A, Tamayo P, Mootha VK, Mukherjee S, Ebert BL. Gene set enrichment analysis: a knowledge-based approach for interpreting genome-wide expression profiles. 2005 Oct 25;102(43):15545–50.

15. Kanehisa M, Goto S, Sato Y, Furumichi M, Tanabe M. KEGG for integration and interpretation of large-scale molecular data sets. 2012 Jan;40(Database issue):D109–14.

16. Ogata H, Goto S, Sato K, Fujibuchi W, Bono H, Kanehisa M. KEGG: Kyoto Encyclopedia of Genes and Genomes. 1999 Jan 1;27(1):29–34.

17. Durinck S, Spellman PT, Birney E, Huber W. Mapping identifiers for the integration of genomic datasets with the R/ Bioconductor package biomaRt. Nat Protoc. 2009;

18. Shen L. GeneOverlap: An R package to test and visualize gene overlaps [Internet]. Mount Sinai; 2013 [cited 2017 Jan 25]. Available from: http://shenlab-sinai.github.io/shenlab-sinai/

19. Gupta S, Ellis SE, Ashar FN, Moes A, Bader JS, Zhan J, et al. Transcriptome analysis reveals dysregulation of innate immune response genes and neuronal activity-dependent genes in autism. Nat Commun [Internet]. 2014;5:5748. Available from: http://www.nature.com/doifinder/10.1038/ncomms6748

20. Parikshak NN, Swarup V, Belgard TG, Irimia M, Ramaswami G, Gandal MJ, et al. Genome-wide changes in lncRNA, splicing, and regional gene expression patterns in autism. Nature [Internet]. 2016 Dec 5 [cited 2017 Jul 18];540(7633):423–7. Available from: www.ncbi.nlm.nih.gov/pubmed/27919067 http://www.nature.com/nature/journal/vaop/ncurrent/pdf/nature20612.pdf

21. Gandal MJ, Zhang P, Hadjimichael E, Walker RL, Chen C, Liu S, et al. Transcriptome-wide isoform-level dysregulation in ASD, schizophrenia, and bipolar disorder. Science (80-) [Internet]. 2018 Dec 14 [cited 2018 Dec 18];362(6420):eaat8127. Available from: http://www.ncbi.nlm.nih.gov/pubmed/30545856

22. Gandal MJ, Haney J, Parikshak N, Leppa V, Horvath S, Geschwind DH. Shared molecular neuropathology across major psychiatric disorders parallels polygenic overlap. Science (80-) [Internet]. 2018;693(February):693–7. Available from: http://biorxiv.org/content/early/2016/02/18/040022.abstract

23. Pulikkan J, Mazumder A, Grace T. Role of the Gut Microbiome in Autism Spectrum Disorders. In: Advances in Experimental Medicine and Biology [Internet]. Springer New York LLC; 2019 [cited 2021 May 25]. p. 253–69. Available from: https://pubmed.ncbi.nlm.nih.gov/30747427/

